# Publishing descriptions of non-public clinical datasets: guidance for researchers, repositories, editors and funding organisations

**DOI:** 10.1101/021667

**Authors:** Iain Hrynaszkiewicz, Varsha Khodiyar, Andrew L Hufton, Susanna-Assunta Sansone

## Abstract

Sharing of experimental clinical research data usually happens between individuals or research groups rather than via public repositories, in part due to the need to protect research participant privacy. This approach to data sharing makes it difficult to connect journal articles with their underlying datasets and is often insufficient for ensuring access to data in the long term. Voluntary data sharing services such as the Yale Open Data Access (YODA) and Clinical Study Data Request (CSDR) projects have increased accessibility to clinical datasets for secondary uses while protecting patient privacy and the legitimacy of secondary analyses but these resources are generally disconnected from journal articles – where researchers typically search for reliable information to inform future research. New scholarly journal and article types dedicated to increasing accessibility of research data have emerged in recent years and, in general, journals are developing stronger links with data repositories. There is a need for increased collaboration between journals, data repositories, researchers, funders, and voluntary data sharing services to increase the visibility and reliability of clinical research. We propose changes to the format and peer-review process for journal articles to more robustly link them to data that are only available on request. We also propose additional features for data repositories to better accommodate non-public clinical datasets, including Data Use Agreements (DUAs).

## Introduction

Open access to research data that can be understood and reused by others is a means to further scientific progress, and publish more reliable and reproducible research^1, 2^. However, clinical research data often include information that could potentially identify individuals, meaning datasets must be anonymised prior to being shared beyond the study for which the data were originally collected. Although guidelines and processes for anonymisation of clinical data exist,^3, 4^ publication of freely available clinical datasets (such as^5^) remains uncommon. As open access to clinical datasets is often unfeasible, a more felicitous and pragmatic approach may be needed.

Some clinical datasets may be made available on request from authors of journal articles or through the recent emergence of dedicated data request systems. However, as large amounts of clinical research can go unpublished^6, 7^, and clinical trials unregistered^8^, the discoverability of many clinical datasets is suboptimal. In this paper we use the term “non-public clinical datasets” to mean datasets that have been generated through experimental clinical research, such as clinical trials, and which are not openly accessible, but are available on request. Clinical research involving surgical, disease-specific or epidemiologic cohort databases, and electronic health records, which can be continually updated and held by an institution, should not be excluded from data sharing but may require specific additional guidance not covered in this paper.

In consultation with relevant stakeholders representing pharmaceutical companies, research funders, researchers and data repository managers, the editors and publishers of the journal *Scientific Data* propose guidelines for linking peer-reviewed journal articles to non-public clinical datasets.

Data repositories are recognised as essential for enabling reliable access to data underlying research in a number of life science disciplines. Their use is ingrained in the practices of some research communities and in the editorial policies of, for example, the Nature journals [Ref 2]. Common repositories for non-public clinical data are less well established, however. We propose implementation of specific features in data repositories for the archival of non-public clinical datasets, to enable appropriate cross-linking of these datasets to scholarly journal articles.

### Summary of recommendations

#### Clinical researchers and their sponsors

- Be prepared to share experimental data with editors, peer reviewers and other researchers in accordance with journal policies
- Apply the shortest possible embargoes on data

#### Repositories

- Develop mechanisms to host clinical research data, including:
  - Provide stable identifiers for metadata records about non-public dataset(s)
  - Implement Data Use Agreements (DUAs)
  - Implement a transparent system for requesting access to data and reviewing requests to access data
  - Allow access to data in a timely manner and include a proportionate review of the scientific rationale, without introducing unnecessary barriers

#### Clinical journal editors and publishers

- Check compliance with their journal’s data sharing policies for every submission
- For manuscripts based on secondary access to trial data (e.g. data the original trial sponsor has made available for further research), check the research is consistent with the DUA and purpose for which data access was granted or ask authors to attest that the submission is compliant with these conditions
- Increase the visibility of clinical datasets to peer reviewers
- Build relationships with repositories for non-public clinical datasets to support public archiving of metadata and data from clinical research
- Introduce links to data, data sharing statements and transparency statements in published articles

#### Data journal editors and publishers

- Develop data article formats (for example Scientific Data’s Data Descriptor) to permanently link articles to descriptions of non-public clinical datasets

#### All sponsors and funders of trials

- Build partnerships with and between data sharing initiatives, trusted repositories, and peer reviewed journals committed to data sharing
- Apply the shortest possible embargoes on data and ensure that data access is subject to a proportionate review (e.g. of the scientific rationale and qualifications of the research team), without introducing unnecessary barriers

## How do researchers currently access non-public clinical data?

Data sharing between researchers has traditionally occurred through direct contact between individuals and research groups^9^. Many journals have policies that require authors to share data that support their results with other researchers on request, but enforcement of such policies varies between journals (for a summary of journal policy types and approaches see^10^). Journal policies that only require data to be “available on request” without also mandating data availability statements, have been found to be less effective for ensuring data access for future researchers^11–14^. However, even clinical journals with strong and enforced policies on data access, (such as the *BMJ* and *PLOS Medicine*), data about identifiable human subjects will usually not be in the public domain, due to the need to protect research participants” privacy.

Alongside changes in journal policies, initiatives from other stakeholders (regulatory agencies, the pharmaceutical industry and research groups) in clinical research have begun to facilitate greater access to non-public clinical datasets. This includes the European Medicines Agency (EMA) which has committed to providing access to individual patient data (IPD) in the future^15^.

When surveyed about their data sharing attitudes and behaviours, clinical researchers express concerns about inappropriate secondary analysis of their data and patient privacy^16^. The Yale Open Data Access (YODA; http://yoda.yale.edu/) project and Clinical Study Data Request (CSDR; http://clinicalstudvdatarequest.com) have since 2012, provided restricted access to non-public clinical datasets while addressing these two concerns. As of January 2016, CSDR listed more than 2800 clinical studies from 12 pharmaceutical companies, for whic data access for future researchershaccess to data could be requested. Researchers are also able to enquire about the availability of other non-listed studies. The project has been described as a success by its independent data access review panel^17^ and 179 research proposals (data requests) were submitted between May 2013 and November 2015 (https://www.clinicalstudydatarequest.com/Metrics.aspx). As of March 2015 GlaxoSmithKline, one of the companies using CSDR, had received 99 requests (approximately four requests per month) to access data from the 1200 trials it had listed. The YODA Project has compiled data from more than 120 studies, representing two commercial data providers, and has received 29 requests (http://voda.yale.edu/table-2-data-requests-submitted).

## Connecting non-public clinical data with journals and repositories

With increasing numbers of open access repositories for research data for many scientific disciplines (http://www.nature.com/sdata/data-policies/repositories), which provide persistent, citable links to datasets and metadata records, it is relatively easy to link journal articles to publicly accessible data. In the area of clinical trials, the concept of allowing secondary researchers to access the data is relatively new and so data are not yet widely shared. In addition, since non-public clinical datasets often have no permanent public record or identifier, links between these and the peer-reviewed literature are far less robust than links between public datasets and the literature. This could be addressed by developing both the data access policies of journals and the relationships of journals with data repositories that can archive non-public datasets. Increasing the visibility of such data could also be supported by a new type of journal article focused on describing non-public clinical datasets – particularly those that are previously unpublished. These new articles could be an adaptation of “data papers” that are already published in data journals. Data journals are a relatively new type of journal focused on data publication^18^. Below we describe the benefits of this approach for researchers, as opposed to simply posting information about non-public datasets on a website or posting summary results of clinical trials in trial registration databases.

### Discoverability

YODA and CSDR provide public information about clinical studies for which data are available on request. Some of these studies may not be described in journal articles and for studies that *are* published it may not be clear in the published article that the underlying data are available. Moreover, when planning a new research project or systematic review, medical researchers primarily rely on bibliographic databases (such as PubMed/MEDLINE, Scopus, Web of Science), individual journals and clinical trial registries to find information rather than these websites. Better links with journal articles, which are more visible due to indexing in bibliographic databases, might increase the visibility and use of data on request services.

*Scientific Data* (http://www.nature.com/scientificdata) from the Nature Publishing Group, is one example of a data journal (see^18^ for others). It is an open access journal for descriptions of scientifically valuable datasets, where data articles (termed Data Descriptors) are linked to their corresponding publicly-available datasets. With non-public datasets journal articles need to link to persistent and citable metadata, a “stub” record or landing page, for a clinical dataset that is available on request. The landing page for a non-public clinical dataset should contain sufficient metadata to facilitate an understanding of what the dataset is, and importantly, the conditions that must be met for access to the data (see below). Data Descriptors (and other types of research-based article) linked to the citeable landing page could then be written by those who created the nonpublic clinical dataset, increasing the discoverability of the dataset and providing a means to obtain scholarly credit for generating and sharing the data.

As well as increasing discoverability of content through indexing in bibliographic databases open access journals, in particular, can help ensure content is highly visible through exposure to standard internet search engines such as Google.

### Quality and peer review

Peer review is central to publishing research in journals. Publication of Data Descriptors at *Scientific Data* (and some other data journals), involves formal peer review by independently-selected experts, of both the article describing the dataset(s) and the dataset itself. The peer-review process at *Scientific Data* focuses on (re)usability and data integrity, rather than on the perceived importance or impact of the data (http://www.nature.com/sdata/for-referees). However, systematic peer review of underlying data is not routine in traditional research journals.

### Data curation

*Scientific Data’s* publication process includes data curation by a dedicated Editor, in addition to peer reviewer checks. This process includes the creation of standardised, machine readable metadata for every Data Descriptor. This is intended to facilitate data discoverability and reuse by using controlled vocabulary terms to capture sample and subject provenance, and outline the experimental workflow (http://www.nature.com/sdata/about/faq#ql4). Datasets in curated, readily consumable formats should increase confidence in the delivered data formats, adding further value to the offerings of data journals. Use of subject-specific repositories, which tend to use community-accepted data formats and standards, and have standardised data curation processes, also increases the likelihood that archived data can be reused.

### Permanence

A role of journals is to ensure the permanence and integrity of the scientific record, for example by maintaining persistent links between articles and datasets, and placing copies of content in redundant archives. Web link decay or “link rot” is well documented, and even in the peer-reviewed literature, an estimated 20% of articles published in 2012 already suffer from broken web links when regular websites or URLs are cited^19^. This deterioration of referential integrity across corresponding data sources presents obstacles to replicating or re-analysing results underlying the scientific record. Publishers’ use of persistent identifiers for journal articles, Digital Object Identifiers (DOIs), helps ensure readers and future researchers can access content as publishers commit to keeping article links up-to-date using the DOI system. DOIs are increasingly being created for datasets and metadata records, via data repositories.

### Links with data repositories

While the YODA Project and CSDR have succeeded in increasing access to clinical research data, their websites, and documents hosted therein are not citable and linkable in the same way as journal articles, and other research objects assigned DOIs. Furthermore, researchers and projects funded by organisations such as Cancer Research UK, Medical Research Council Clinical Trials Unit (MRC CTU)^20^ and the Wellcome Trust may have well-managed non-public datasets available via local hosting, but these archives are unlikely to meet preservation, discoverability and linking standards of journals and publishers. *Scientific Data*, for example, works with trusted repositories to publish its Data Descriptors and supports data archiving policies and activities across the *Nature* journals^2^. *Scientific Data* has established criteria for assessing public research data repositories (see below) and has so far approved more than 80 repositories (http://www.nature.com/sdata/data-policies/repositories; https://biosharing.org/collection/ScientificData). Other publishers and journals also list suggested or recommended repositories for authors including BMJ Open, PLOS, BioMed Central.

### Negative, incomplete or inconclusive trial data

While a number of journals exist that explicitly encourage negative or inclusive trials to be published (such as *Trials, Journal of Negative Results in Biomedicine*), a type of article designed to describe non-public clinical trial data – regardless of whether discussion or analysis of the data are available – might be further incentive for researchers and their sponsors to share data.

## Additional considerations for journals and data repositories

Data repositories that host non-public clinical datasets which are linked to peer-reviewed articles, need to provide the following additional services to those repositories hosting only publicly-accessible data.

### Data use agreements

An essential component for the secondary use of non-public clinical datasets is a data use agreement (DUA) between the data generator or repository and the secondary researcher(s). The purposes of DU As include reducing risks to participants and other parties involved in the study and to ensure the scientific value of secondary analyses, which includes commitments to publishing secondary analyses in peer-reviewed journals^21^.

Template DUAs are provided by the YODA Project and CSDR (https://www.clinicalstudvdatarequest.com/Documents/DATA-SHARING-AGREEMENT.pdf). The Multi-regional Clinical Trials Center at Harvard University also provides a template DUA (http://mrct.globalhealth.harvard.edu/file/376776).

Recommending wording for DUAs is beyond the scope of these guidelines. However, the position of *Scientific Data* is that DUAs describing datasets intended for publication in journals dedicated to open science should support wider use by independent qualified researchers, including for competitive analysis, and that the authors agree to relinquish intellectual property claims stemming from the data. Mandatory requirements for co-authorship for data creators should be avoided.

### Controlled access and governance

Few repositories have processes for managing and approving requests to access non-public clinical datasets but we identify some candidates and possible candidates below (Table 1). Controlled access approaches for IPD have been described by Tudur Smith *et al*.^22^ – which, for clinical trials, do not always use data repositories – can include independent advisory or data access committees. The role of a trusted intermediary for repositories for clinical data is also highlighted by the IOM’s report^21^ but there is no widely established solution for all experimental clinical research. Some concerns have been raised about the speed of data release and usability of data from “voluntary data-sharing portals”^23^ such as YODA and CSDR and, more broadly, clinical trial units are understandably concerned about costs of managing controlled access to data^24^.

**Table 1.** Data repositories that meet or potentially could meet the proposed requirements (in this article) for hosting non-public clinical trial data.

| **Repository Name** | **Repository URL** | **Type of data hosted** | **Access controls** | **Example of non-public clinical dataset** |
| --- | --- | --- | --- | --- |
| UK Data Service - ReShare | http://reshare.ukdataservice.ac.uk | All | Requires account to request access to specific dataset | https://dx.doi.org/10.5255/UKDA-SN-851861 |
| ICPSR | https://www.icpsr.umich.edu | All | Requires account to request access to specific dataset | http://doi.org/10.3886/ICPSR23580.v2 |
| European Genome-phenome Archive (EGA) | https://www.ebi.ac.uk/ega | genomics data | Specific to each study, some data are open | https://www.ebi.ac.uk/ega/datasets/EGAD00000000031 |
| database of Genotypes and Phenotypes (dbGaP) | http://www.ncbi.nlm.nih.gov/gap | human genotype and phenotype data | Requires account to request access to specific dataset | http://www.ncbi.nlm.nih.gov/projects/gap/cgi-bin/study.cgi?study_id=phs000001 |
| Harvard Dataverse | http://dataverse.harvard.edu | All | Specific to each study, some data are open | http://dx.doi.org/10.7910/DVN/25833 |
| figshare | http://figshare.com | All | Specific to each study, some data are open | https://dx.doi.org/10.17608/k6.auckland.2003298 |
| National Database for Clinical Trials related to Mental Illness (NDCT) | http://ndct.nimh.nih.gov | NIH funded data on any aspect of human mental health [includes National Database for Autism Research (NDAR) and Research Domain Criteria Database (RDoCdb)] | Requires account to request access to specific dataset | https://ndar.nih.gov/edit_collection.html?id=14 |
| National Addiction & HIV Data Archive Program (NAHDAP) | http://www.icpsr.umich.edu/icpsrweb/NAHDAP/index.jsp | drug addiction and HIV research data | Requires account to request access to specific dataset | http://doi.org/10.3886/ICPSR33581 |
| Cancer Imaging Archive | http://www.cancerimagingarchive.net/ | anatomical imaging of human cancer | Requires completion of a DUA, some data are open | https://wiki.cancerimagingarchive.net/pages/viewpage.action?pageId=20643859 |
| Synapse | http://sagebase.org/synapse | biomedical data | Specific to each study, some data are open | https://dx.doi.org/10.7303/syn4993293 |

### Landing pages

Repository metadata records (landing pages) for linking journal articles to non-public clinical data will share many characteristics with repository records for public datasets. Good practice for data citation already recommends that links to datasets should resolve to landing pages rather than the raw data files, and standards are emerging for the information that is essential and desirable to be included in landing pages^25^. These standards consist of a dataset description comprising a dataset identifier, title and brief description, creator, Publisher, and publication year. Landing pages should also include persistence/permanence information and licensing information for the data. We recommend that landing pages for non-public clinical datasets also include information detailing the access controls pertinent to the data. Examples of landing pages can be found at the UK Data Archive (e.g. http://dx.doi.org/10.5255/UKDA-SN-7612-2) and the European Genome-phenome Archive (e.g. https://www.ebi.ac.uk/ega/datasets/EGAD00000000001). For clinical trials, where prospective trial registration is established as ethical research practice, we recommend unique trial identifiers be included on landing pages.

### Additional repository assessment criteria

Taking the above issues into consideration, we propose additional criteria by which journals and publishers can assess repositories for hosting of non-public clinical datasets:

### Trusted repositories for non-public datasets must

- Provide stable identifiers for metadata records about non-public dataset(s)
- Allow access to data with the minimum of restrictions needed to ensure protection of privacy and appropriateness of secondary analyses, codified in Data Use Agreements (DUAs)
- Allow access to data in a timely manner
- Provide support for users of data

### Trusted repositories for non-public datasets should, ideally*

- Be independent of the study sponsors and principal investigators
- Have a transparent and persistent system for requesting access to data and reviewing requests to access data
- Provide relevant data analysis software environments where data are not permitted to be downloaded locally
- Provide public access to the metadata of archived data for third party search and discovery functionalities

These criteria are in addition to the current repository selection criteria (http://www.nature.com/sdata/about/faq#q21) of *Scientific Data* which require trusted repositories:

- Be broadly supported and recognized within their scientific community
- Ensure long-term persistence and preservation of datasets in their published form
- Implement relevant, community-endorsed reporting requirements
- Provide stable identifiers for submitted datasets
- Allow public access to data without unnecessary restrictions

Since this working group was formed, at least one other group (Burton et al) has proposed criteria for “data safe havens” in healthcare^26^.

### Repositories that could or may meet the current and additional requirements

Specialist trial data systems such as CSDR (and YODA) have indicated a willingness to listen to feedback from researchers and may evolve as data sharing progresses. As well as the potential for these repositories to develop to meet certain criteria, there are number of other general and specialist repositories that could or may meet the above criteria (Table 1), at least for specific types of clinical research.

There are also clinical data systems and “federations” associated with specific projects and some have played roles in disseminating data in specific communities. The ADNI portal is one such example (http://neuroinformatics.harvard.edu/gsp/loni). The group responsible for this portal have published a peer-reviewed article^27^ describing and linking to non-public data in this resource.

Another Data Descriptor published at *Scientific Data* describes a non-public clinical dataset – of neuroimaging data from brain tumour patients – hosted at the UK Data Archive^28^.

### Additional manuscript sections required

The format of published articles – data papers and traditional research articles – needs to be developed to accommodate links to non-public datasets.

Articles should include information about why the data are not publicly available (i.e. because they contain personally identifiable information) and describe the restrictions on accessing the dataset. They should also state if the data are subject to a DUA and where the DUA can be found – ideally, this should be hosted permanently with the landing page of the non-public dataset. In *Scientific Data*, the Usage Notes section would be appropriate for this information. Persistent links to landing pages should also be cited, in the same way public datasets are cited. Other journals, such as *PLOS ONE, Palgrave Communications, GigaScience* and *Royal Society Open Science* are now routinely including dedicated article sections to describe and link to datasets supporting published articles – these could be adapted to meet these requirements.

Where Data Descriptors, and other articles, link to non-public clinical datasets we recommend authors include in their articles a transparency declaration guaranteeing that their description of the dataset is an honest and accurate account. Transparency statements for regular journal articles, for other aspects of research integrity, have been implemented by the BMJ^29^.

See Figure 1 for an overview of the standard editorial workflow of *Scientific Data*, and Figure 2 for the proposed modified workflow to accommodate Data Descriptors of non-public clinical datasets.

**Figure 1.**
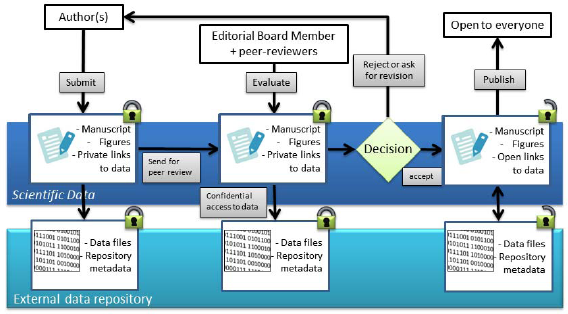
Overview of standard Scientific Data editorial workflow for non-confidential datasets. With submission of a Data Descriptor, authors provide a secure link to dataset(s) stored in an external repository. Editors and referees are granted access to the data, in a manner that does reveal their identities to the authors. Upon publication, both the article and the datasets are made freely accessible online under appropriate terms or licences.

**Figure 2.**
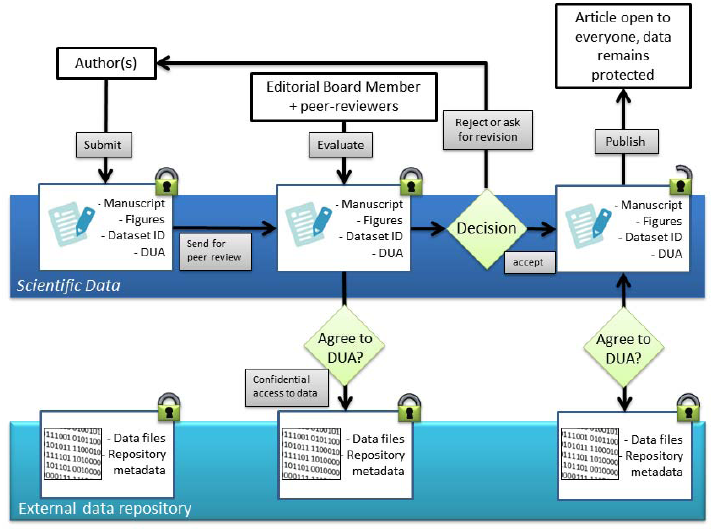
Overview of an example editorial workflow accommodating peer review and publication of clinical Data Descriptors. For Data Descriptors describing a clinical restricted-access dataset, authors provide with their submission a description of where the dataset(s) are hosted and a copy of the DUA. A process is then agreed between the journal and the authors, by which referees and editors may request access to the data during peer-review. Upon publication, the article is made openly available, and the host data repository releases a landing page for the clinical dataset(s). Users may request access to the data according to the process and terms outlined in the Data Descriptor and associated DUA.

### Research participant consent

Part of a journal’s role is to enforce relevant ethical expectations regarding consent. An important consideration is whether participants gave appropriate consent for data to be made available to secondary researchers in the future (if data are not fully anonymised^3^). Where informed consent for data sharing was obtained, consent should include scholarly publications (peer-reviewed articles) that describe or link to datasets.

## What data should be available to secondary researchers?

Different types of experiments produce different types of information in a variety of formats, leading to different minimum requirements for secondary researchers seeking to replicate or understand results. In general, reproducible medical research requires access to data, code, and study protocols^30^. CSDR has, furthermore, defined data and document types for the studies it lists, although all items are not always available for each study:

i. Raw dataset
ii. Blank case report forms
iii. Annotations of blank case report forms
iv. Dataset specifications
v. Protocol (all versions)
vi. Analysis-ready dataset
vii. Reporting and analysis plan
viii. Clinical study report

While the scope of these guidelines potentially goes beyond clinical trials – including molecular data types – this list defined by CSDR is a reasonable guide. The IOM has also described the clinical research data types that are needed for reanalysis, which differs slightly from the CSDR list (Chapter 5, p112)^21^. Standards for reusability for historic, non-public clinical datasets might need to be less stringent if data are only available in file formats that might not be optimised for reuse.

## Who should have secondary access to data?

The consensus of regulators, industry sponsors and funders of clinical trials is for data access to be granted to suitably qualified researchers with a legitimate reanalysis proposal. A requirement of CSDR’s procedures is that a researcher with a degree in statistics or a related discipline should be part of the research team. The Independent Review Panel for CSDR had, as of February 2015, approved 71 requests and rejected or advised to resubmit 4 requests. The YODA Project’s approval committee assesses “basic information about the Principal Investigator, Key Personnel, and the project Research Proposal, including a scientific abstract and research methods” when reviewing data access requests^31^.

Journal polices generally require data supporting submitted works must be accessible to peer reviewers and editors, and study sponsors should already be used to providing access to data supporting manuscripts submitted to major medical journals. These repository criteria and guidelines could make these processes more efficient if applied to clinical research journals. To publish Data Descriptors in *Scientific Data*, peer reviewers and editors must be given controlled access to supporting data for every article. The majority of journals operate single or double blind peer review, which means some reviewers or journals might require their anonymity to be maintained (public data repositories often support anonymous peer review, although there is increasing adoption of open peer review).

## Peer review of non-public clinical datasets

Journals that implement these guidelines will be able to make non-public clinical datasets more visible to peer reviewers, potentially, as well as editors. *Scientific Data* is reviewing its peer review guidelines (http://www.nature.com/sdata/for-referees#writing-review) as it considers the first articles describing non-public datasets but many journals have their own guidelines and processes for their peer reviewers. In general, however, peer review of articles describing and linking to nonpublic clinical datasets should include but not be limited to:

- Whether the access controls on the data are warranted and if enough information is provided on how to request access to the data
- Whether the data are sufficiently well described to enable independent researchers to assess the reuse value of the dataset (to help ensure data requested are reused)

## When to provide secondary access to data

The IOM has recommended embargoes of up to 18 months from study completion before clinical trialists are required to share data, although this has been criticised for being too long^32^. In a major epidemic, a long embargo on data access and reuse could be to the detriment of fighting disease. But a reasonable period for analysis – a right of first use – is acknowledged in most research communities. Any embargo on non-public clinical dataset(s) described in and linked to a journal article would have to have expired to comply with the recommendations in this article. In general we advocate no, or short, embargoes on data release wherever feasible.

## Next steps

In consultation with a working group (see Acknowledgements) *Scientific Data* is developing its editorial and peer-review processes and relationships with repositories to support publication of Data Descriptors for non-public clinical datasets as we receive relevant submissions. Other journals – data journals and traditional journals – may wish to consider these repository, linking and editorial policy proposals. Some members of our working group are also helping to identify interested research teams and relevant datasets that could be part of a publication pilot. Indeed, we need real data with which to develop more robust links between non-public datasets and journal articles. We strongly encourage others to contact the editors, to discuss proposals.

## Acknowledgements

Thanks to the Working Group members, who attended a meeting in December in 2014 which identified many of the issues highlighted in this paper and who provided comments on earlier drafts of the paper:

Robert Frost, Policy Director, Medical Policy, GSK
Richard Lehman, on behalf of the Yale Open Data Access Project
Fiona Reddington, Head of Clinical and Population Research Funding, Cancer Research UK
Will Greenacre, Policy Officer, Wellcome Trust
Matthew Sydes, Senior Scientist and Statistician, MRC Clinical Trials Unit
John Gonzalez, Publications Director, AstraZeneca
Jay Bergeron, Director, Translational and Bioinformatics, Pfizer
Jessica Ritchie/Joseph Ross, Yale Open Data Access Project
Mark Hahnel, CEO and Founder, figshare

Thanks also to those who submitted comments on the first public version of these guidelines, which was announced in July 2015^33^’^34^.

